# Perfusion quality does not necessarily predict ultrastructural preservation after hyperosmotic brain perfusion

**DOI:** 10.64898/2026.07.27.741074

**Authors:** Andria Slaughter, Macy Garrood, Katelyn Hedden, Allison Sowa, William Janssen, Emma L. Thorn, Claudia De Sanctis, Kurt Farrell, John F. Crary, Andrew T. McKenzie

## Abstract

Perfusion fixation is widely used in neuroscience to prepare mammalian brain tissue for histological and ultrastructural analysis. Perfusion protocols are commonly assessed using macroscopic indicators such as gross appearance and neuroimaging, which assess the extent to which perfusate has been distributed throughout the brain. There is a critical need to determine to what extent these metrics can accurately predict high-quality ultrastructural preservation, particularly as new perfusion protocols are developed for connectomics. In this technical report, we describe evidence that these two measures can be decoupled by the addition of dehydrating agents to the perfusate solution. In three human brain donors and one canine brain donor perfused with a fixative solution containing 10% mannitol and 10% polyethylene glycol 35 kDa, macroscopic and radiological indicators of perfusion quality appeared adequate or favorable. However, electron microscopy revealed expanded extracellular space, shrunken cellular processes, and distorted cell membranes, consistent with an osmotic shock artifact resulting from severe hyperosmotic dehydration. Similar ultrastructural artifacts were observed in a canine brain donor perfused with 20% mannitol in 20% neutral buffered formalin without PEG. We compare these ultrastructural findings with findings from previously reported cases perfused with standard neutral buffered formalin without osmotic additives. These findings illustrate a risk of optimizing brain perfusion protocols designed to preserve neural circuitry based on macroscopic or radiological perfusion quality metrics alone, since these metrics can be satisfied while the ultrastructure is severely compromised.

## Introduction

The artificial perfusion of fixative solutions through the cerebral vasculature has long been the standard method for preserving mammalian brain tissue for histological and ultrastructural analysis [1–3]. The approach is widely used because perfusion can distribute fixative solutions much faster across the brain, thereby reducing artifacts due to autolysis. This technique appears to be essential for preparing whole mammalian brains for connectome imaging with contemporary methods, which is an aspirational goal of the connectomics community [4,5]. The quality of perfusion can be assessed using macroscopic and radiological indicators, including visual inspection of vascular clearance, tissue color changes indicating fixative penetration, and neuroimaging modalities such as computed tomography (CT) that can reveal the spatial distribution of perfusate throughout the brain parenchyma [6].

A key challenge in perfusion-based brain preservation is that even relatively short periods of ischemia can progressively impair subsequent perfusion [7,8]. One strategy to overcome this is to include osmotically active solutes in the perfusate. Because the blood-brain barrier limits the rate at which many solutes equilibrate with brain tissue, these additives can establish osmotic or oncotic gradients that draw water out of the parenchyma, reducing perivascular edema and potentially improving vascular perfusion [9,10]. The duration and mechanism of these gradients depend on the properties of the solute. Perfusate additives that cannot readily cross the BBB, such as high-molecular-weight polyethylene glycols (PEGs), generate sustained oncotic gradients that draw water out of the brain parenchyma, thus relieving or preventing perivascular edema and potentially improving perfusion following a period of ischemia [11,12]. Smaller solutes such as mannitol, which cross the BBB slowly relative to water, can generate transient osmotic gradients during perfusion that can improve flow through similar mechanisms.

However, perfusion with additives that generate these osmotic gradients can also cause hyperosmotic dehydration of the tissue. A significant problem in severe cases of such hyperosmotic dehydration is osmotic shock artifact, which indicates the distortion or disruption of the fine processes, synaptic structures, and other cellular and extracellular structures that constitute the brain’s ultrastructure, as a result of rapid water shifts [9,13]. Similar concerns have been raised regarding cryoprotective agent perfusion, wherein a dehydration-induced ultrastructural artifact has been recognized as a potential obstacle to tracing fine neural processes, motivating the use of perfusion fixation prior to cryoprotectant loading [14].

At typical concentrations, the aldehyde fixative itself contributes relatively little to the effective osmotic concentration that mediates the extent to which cells shrink or swell [15–17]. This is because aldehydes cross cell membranes freely, so they raise the total measured osmolarity of a solution without raising its effective osmotic concentration. Instead, effective osmolarity is determined by the non-aldehyde fraction of the perfusate (i.e. the carrier solution), including the buffer salts and any added solutes. In rat brain perfusion, it has been reported that structural preservation quality is best when the fixative buffer is present at near or slightly above standard tissue tonicity [18]. On the other hand, when a high osmotic concentration of the fixative carrier solution has been used, the reported effect is shrunken cells and processes with expanded extracellular space, albeit with preserved myelin periodicity [18]. Taken together, the practical implication is that the composition of the carrier solution, rather than the aldehyde itself, is the variable that determines the osmotic effects of perfusion fixation in the brain.

Despite these theoretical concerns, there is not a substantial amount of data comparing perfusion quality metrics with ultrastructural outcomes when dehydrating agents are incorporated into brain perfusates. Here, we present data from two brain banking experiments addressing this question. In the first experiment, we evaluated four donors (three human, one canine) perfused with a fixative solution containing 10% mannitol and 10% polyethylene glycol 35 kDa (PEG35). In the second experiment, we evaluated a canine donor perfused with 20% mannitol in 20% neutral buffered formalin (NBF) without PEG. In both experiments, we found that the addition of these agents appeared to cause severe ultrastructural artifacts visible on electron microscopy.

## Methods

### Brain banking procedures

Anatomical whole-body donations were performed by a partner whole-body donation organization operating under Oregon Health Authority regulations, as previously described [19]. Additionally, through our canine brain bank program, we were donated the bodies of two deceased canines following euthanasia by a licensed veterinarian, with signed owner consent for research use [20]. The Apex Neuroscience Brain and Tissue Bank operates under an exemption determination issued by the Pearl Institutional Review Board (IRB) after the submission of our protocols for review.

### Brain tissue perfusion methods

Human brains were perfused through the bilateral common carotids or internal carotid arteries using our previously described methods [6]. Canine brains were perfused after transcardial or aortic cannulation using our previously described methods [20]. In both species, the base perfusate was NBF, with perfusate composition varying by the particular case under consideration (**Table 1**). Concentrations of mannitol and PEG35 are reported as weight per volume (w/v). Iohexol (3 mgI/mL) was included in most cases to facilitate visualization of perfusate distribution on post-perfusion CT imaging. A small amount of colored dye was also added to aid in the visual assessment of perfusate distribution. When available, post-perfusion CT imaging was performed using a 16-slice OmniTom Elite scanner (Neurologica, Danvers, MA). Perfusion quality was evaluated qualitatively by gross examination and CT imaging using previously described assessment methods based on tissue appearance and the spatial distribution of iohexol contrast [6]. Note that donors 7, 34, and 65 were previously reported cases [19] perfused without osmotic additives that are included here for comparison.

**Table 1.** Characteristics of brain donors included in this study. For donor 327, the precise time of death was not recorded, but the PMI is known to be less than one hour. *: This donor utilized medical aid in dying (MAID) for end-of-life care. **: For this donor (but none of the others), 1-1.5 L of washout solution consisting of 1x phosphate buffered saline, iohexol, and a small amount of blue dye was perfused prior to the perfusion of fixative solution. PMI: Postmortem interval. NBF: Neutral Buffered Formalin. PEG35: Polyethylene glycol 35 kDa. NR: Not recorded.

| Donor ID | Age | Sex | PMI | Reported cause of death | Fixative perfused | Fixative perfusate additives | Fixative volume perfused | Duration of perfusion | Perfusion route |
| --- | --- | --- | --- | --- | --- | --- | --- | --- | --- |
| 7 | 78 | M | 4.25 hr | Cancer* | 10% NBF | None** | 3L | 15 min | Common carotids |
| 34 | 88 | M | 20 hr | Cancer* | 10% NBF | Green dye | 3L | 20 min | Common carotids |
| 65 | 17 | F | 1.5 hr | Euthanasia (Donated canine) | 10% NBF | None | NR | NR | Transcardial |
| 247 | 79 | M | 14 hr | Cancer* | 10% NBF | **10% mannitol, 10% PEG35, iohexol, green dye** | 2L | 17 min | Internal carotids |
| 250 | 15 | M | 43 min | Euthanasia (Donated canine) | 10% NBF | **10% mannitol, 10% PEG35, iohexol, green dye** | NR | 23 min | Aorta |
| 252 | 87 | M | 24 hr | Chronic Heart Failure | 10% NBF | **10% mannitol, 10% PEG35, iohexol, green dye** | 2L | 31 min | Internal carotids |
| 256 | 69 | F | 16 hr | Chronic Heart Failure | 10% NBF | **10% mannitol, 10% PEG35, iohexol, green dye** | 5L | 54 min | Internal carotids |
| 327 | 13 | M | <1 hr | Euthanasia (Donated canine) | 20% NBF | **20% mannitol, iohexol, green dye** | 8L | 25 min | Aorta |

### Osmolality testing

Perfusate osmolality was measured by freezing point depression using an OsmoTECH XT Single-Sample Micro-Osmometer (Advanced Instruments, Norwood, MA, USA). The mean measured osmolality for 10% NBF was 1811 mOsm/kg H_2_O (CV = 0.26%, n = 5), for 10% NBF with 10% mannitol and 10% PEG35 was 2764 mOsm/kg H_2_O (CV = 0.38%, n = 5), for 20% NBF was 3049 mOsm/kg H_2_O (CV = 0.46%, n = 5), and for 20% NBF with 20% mannitol was 4055 mOsm/kg H_2_O (CV = 0.75%, n = 5).

### Light microscopy

Brain tissue from the frontal pole and adjacent white matter was placed into a tissue cassette for processing and paraffin embedding. Sections (6 μm thick) were cut from the paraffin-embedded tissue, baked, deparaffinized, and stained with hematoxylin and eosin (H&E). Whole-slide images (WSIs) were acquired at 40x using an Aperio GT450 slide scanner (Leica Biosystems).

### Electron microscopy

Tissue samples from selected brain regions were processed for electron microscopy. For donors 247, 250, 252, and 256, samples were examined from the frontal cortex, adjacent frontal cortex white matter, and thalamus. For donor 327, samples were examined from the bilateral cerebellum, bilateral frontal cortex, bilateral temporal cortex, thalamus, and temporal white matter. Within cortical samples, imaging was performed primarily in layers II/III.

Briefly, tissue was post-fixed and processed using an adapted NCMIR heavy-metal staining protocol [21]. This included sequential treatments with tannic acid, reduced osmium, thiocarbohydrazide, osmium, and uranyl acetate at room temperature, followed by lead aspartate staining at 60°C. Samples were dehydrated through graded ethanol, infiltrated with Embed 812 epoxy resin (EMS), and polymerized for 72 hours at 60°C. Ultrathin sections (70 nm) were cut using a Leica UC7 ultramicrotome and collected on nickel slot grids. Images were acquired on two systems. First, a HT7500 transmission electron microscope (Hitachi High-Technologies, Tokyo, Japan) using an AMT NanoSprint12 12-megapixel CMOS TEM Camera System, with minimal contrast adjustments applied during acquisition. Second, a Crossbeam 350 field-emission scanning electron microscope/focused ion beam microscope (ZEISS, Oberkochen, Germany) equipped with a Gemini field-emission electron column and an annular scanning transmission electron microscopy (aSTEM1) detector. Images were acquired in STEM mode using a 15 µm objective aperture, with accelerating voltage and working distance optimized for each specimen. Prior to imaging on the Crossbeam 350 system, selected STEM grids were sequentially post-stained with uranyl acetate followed by Reynolds’ lead citrate [22]. Grids were floated on a drop of uranyl acetate for 15-30 min at room temperature in the dark, rinsed thoroughly with distilled water, and then floated on Reynolds’ lead citrate for 5-10 min at room temperature before a final rinse with distilled water.

## Results and discussion

### Perfusion quality and preservation artifacts with PEG35

We first evaluated four cases (three human donors and one canine donor) perfused with a fixative solution containing 10% mannitol and 10% PEG35. These additives were included to generate osmotic and oncotic gradients intended to improve perfusion after ischemia. Across all four of these cases, qualitative measures indicated adequate to favorable perfusion quality (**Figure 1**). Gross examination showed largely blood-free surface vessels and a dehydrated appearance on the cortical surface. Notably, these brains were also found to be somewhat more friable during extraction than our previous NBF-only cases. CT demonstrated relatively symmetrical perfusate distribution, with more complete opacification of cortical grey matter than deep white matter, consistent with previous reports that white matter is more difficult to artificially perfuse after ischemia [23]. Light microscopy likewise showed that both large and small vessels in the sampled regions were largely cleared of blood. Overall, the perfusion quality appeared broadly comparable to that observed in our previously reported NBF-only cases.

**Figure 1.**
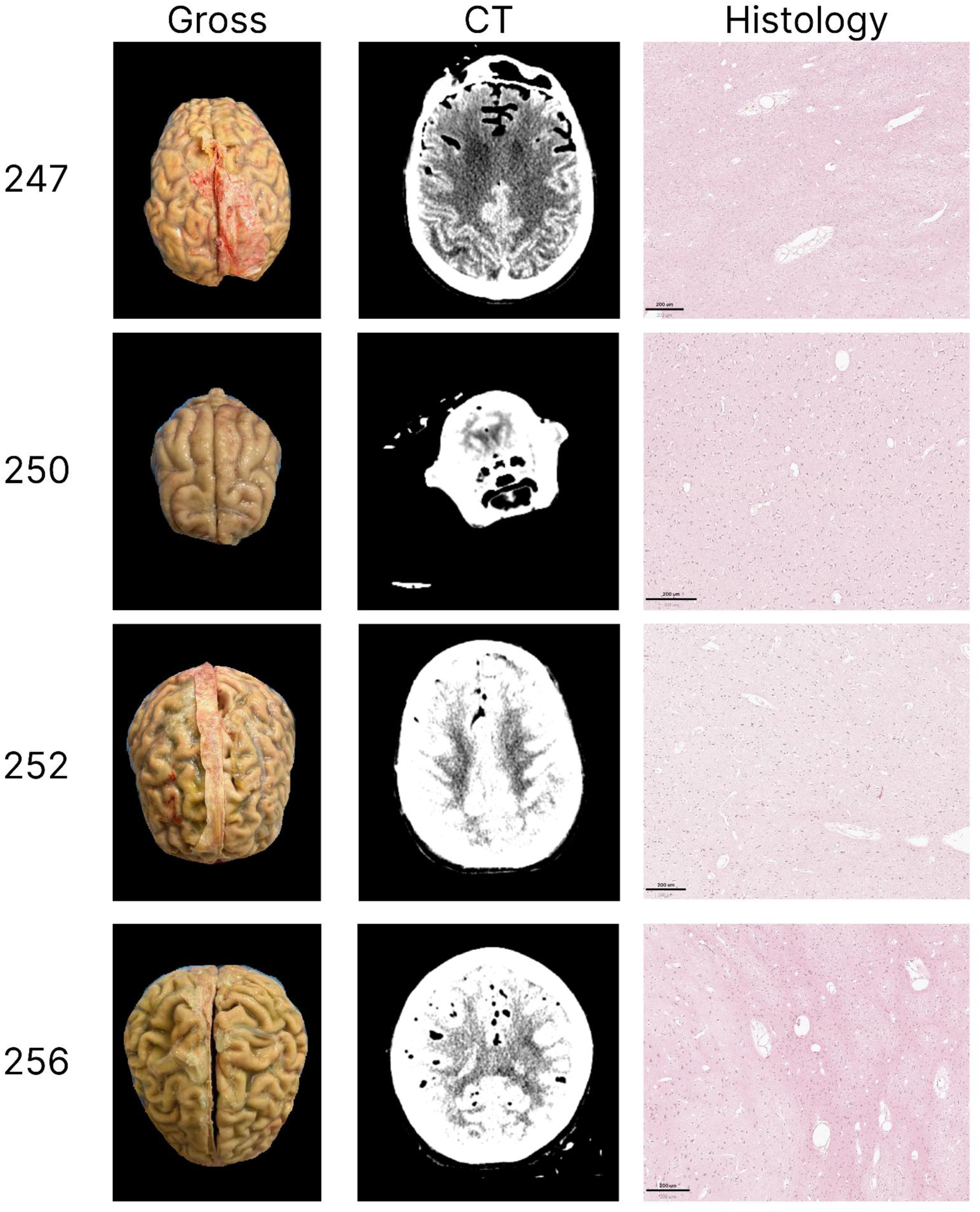
Representative gross, computed tomography (CT), and light microscopic findings following perfusion with 10% NBF, 10% mannitol, and 10% PEG35. Representative images are shown from the brains of four donors (247, 250, 252, and 256), of which 250 is a canine donor and the rest are human donors. Columns show gross appearance, post-perfusion CT, and H&E-stained histological sections from the frontal cortex grey matter. Scale bars for histology images: all 200 µm.

On light microscopy, the frontal cortex and adjacent white matter showed heterogeneous changes that were generally consistent with an artifact caused by severe hyperosmotic dehydration during perfusion (**Figure 2**). In the grey matter, the most characteristic feature of this artifact was a banded pattern in which compact, eosinophilic tissue alternated with pale regions showing apparent expansion of the extracellular space. We also observed cellular shrinkage and pericellular halos, although these can also be seen in tissue with long PMI prior to fixation [24]. The white matter showed patchy rarefaction. In some areas, this was distributed in a banded pattern, alternating with relatively compact tissue. Despite these abnormalities, the overall tissue architecture was still recognizable on light microscopy.

**Figure 2.**
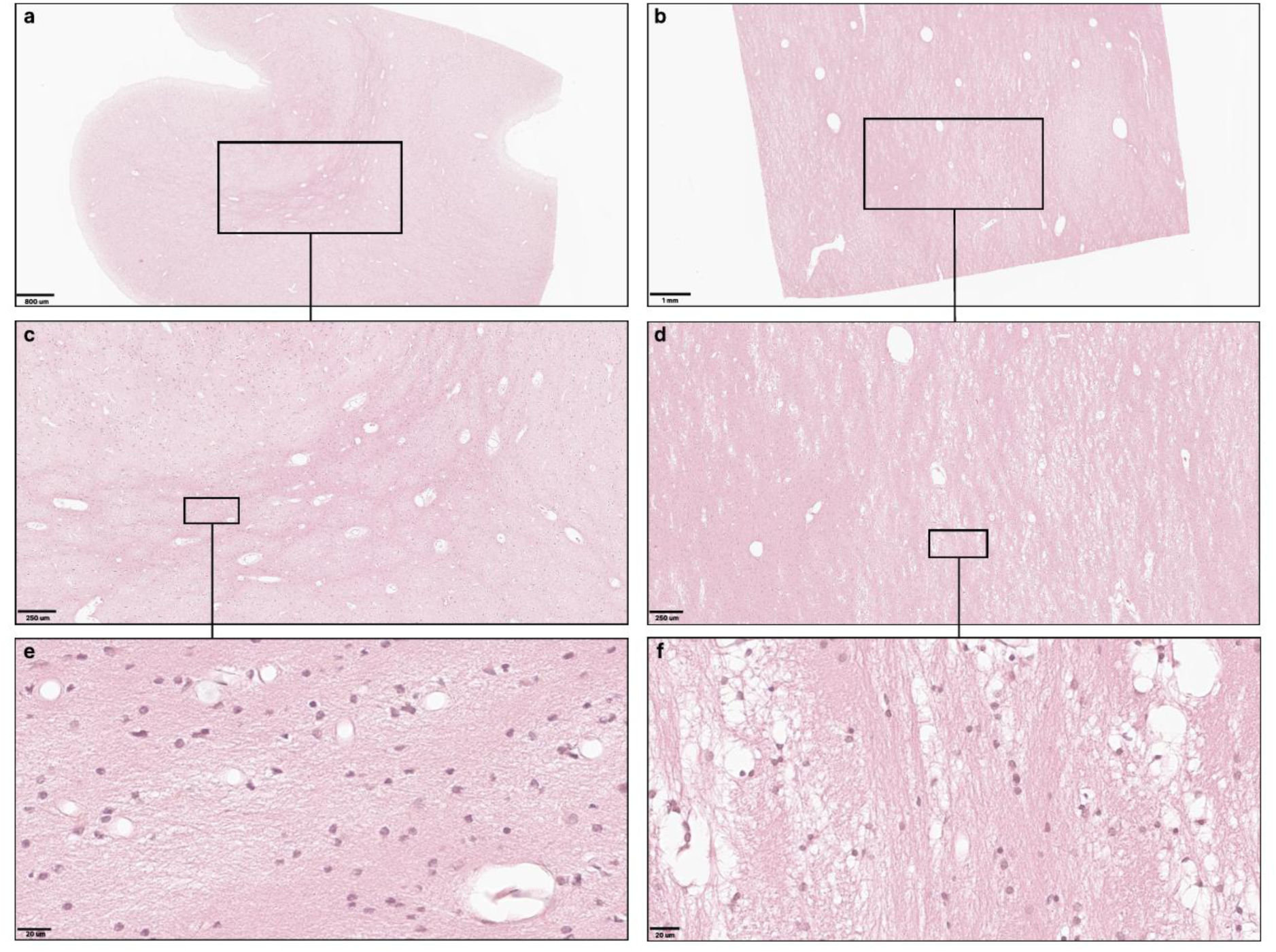
Representative light microscopy findings following perfusion with 10% NBF, 10% mannitol, and 10% PEG35. Images are from donor 256. Panels (**a**), (**c**), and (**e**) show frontal cortex grey matter, and panels (**b**), (**d**), and (**f**) show adjacent white matter. Successive rows show increasing magnification of the boxed regions in the preceding row. Scale bars: (**a**) 800 µm; (**b**) 1 mm; (**c**, **d**) 250 µm; (**e**, **f**) 20 µm.

On electron microscopy, all four brains perfused with 10% mannitol and 10% PEG35 showed severe ultrastructural abnormalities across all three brain regions examined (**Figure 3**). The normally compact neuropil was replaced by widely separated, electron-dense cellular processes surrounded by large electron-lucent spaces. Fine cellular processes were frequently shrunken or fragmented, and cell membranes were often distorted or discontinuous. Myelin sheaths were also frequently abnormal, with thickened, intensely electron-dense, and irregularly folded lamellae rather than the normal concentric organization.

**Figure 3.**
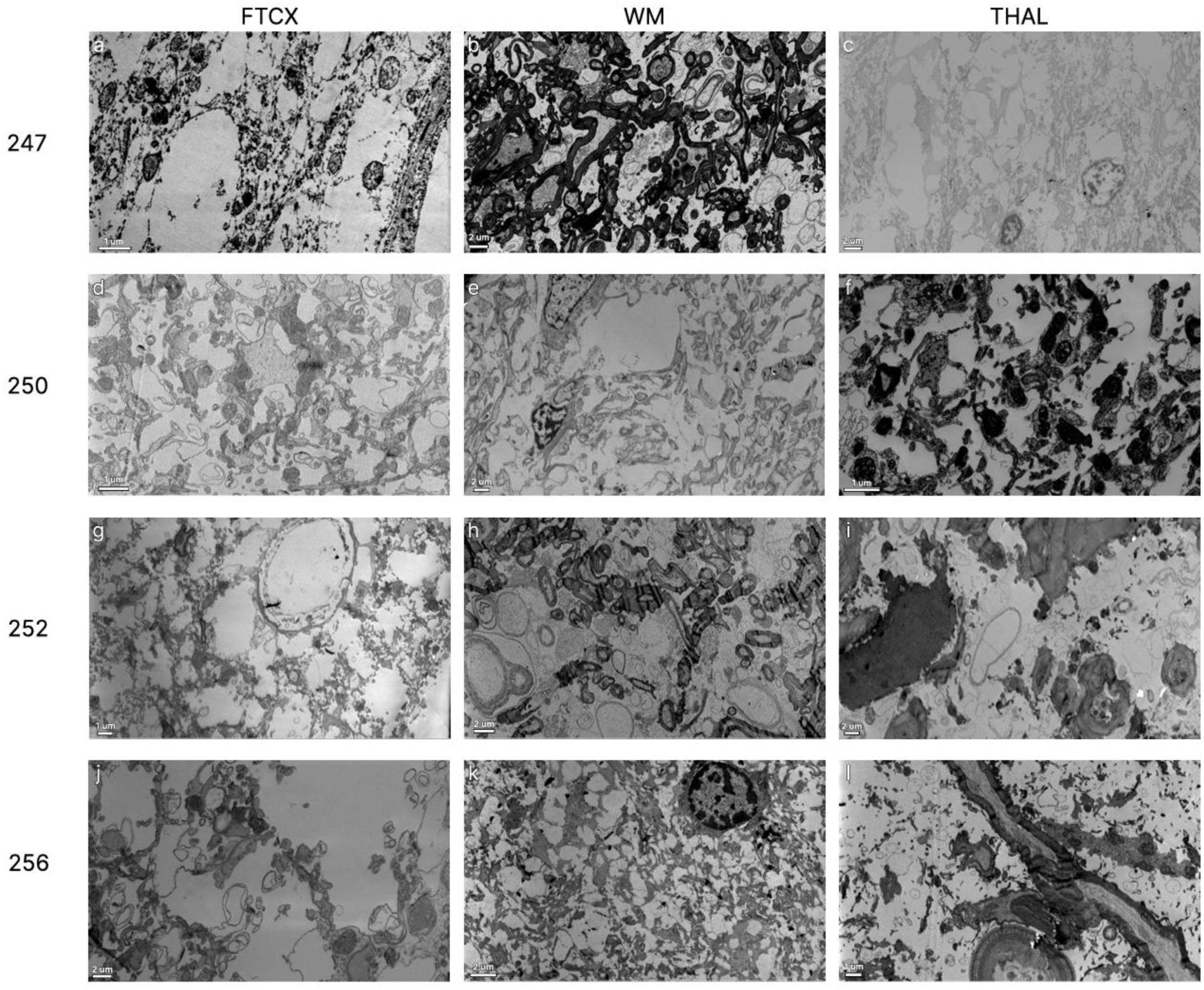
Representative electron micrographs following perfusion with 10% neutral buffered formalin (NBF), 10% mannitol, and 10% PEG35. Representative images are shown from donors 247, 250, 252, and 256, of which 250 is a canine donor and the rest are human donors. Columns correspond to frontal cortex (FTCX), white matter (WM), and thalamus (THAL). Images were acquired using a Zeiss Crossbeam 350 scanning electron microscope in STEM mode. Several samples (**a**, **b**, **e**, **f**, **k**) were post-stained with uranyl acetate and lead prior to imaging. Scale bars: 1 µm (**a**, **d**, **f**, **g**, **l**), 2 µm (**b**, **c**, **e**, **h**, **i**, **j**, **k**).

### Preservation artifacts with high-concentration mannitol

The severe ultrastructural abnormalities observed with 10% mannitol and 10% PEG35 could theoretically have reflected a PEG-specific effect rather than hyperosmotic dehydration itself. To distinguish between these possibilities, we evaluated a canine brain perfused with 20% mannitol in 20% NBF without PEG. In this case, gross examination showed a pale brain with largely blood-free surface vessels and a firm consistency (**Figure 4**). Although these gross findings suggested high-quality perfusion, electron microscopy revealed severe ultrastructural artifacts in all eight brain regions examined (**Figure 5**). These alterations were similar to those observed in the brains perfused with 10% mannitol and 10% PEG35, including marked expansion of the extracellular space, shrunken and distorted neurites, frequent membrane disruption, and irregularly folded myelin sheaths. In regions containing larger cellular profiles, the plasma membranes of the cell body and surrounding pale glial processes were not clearly visible, in contrast to the crisp membrane boundaries typically visible in well-preserved tissue.

**Figure 4.**
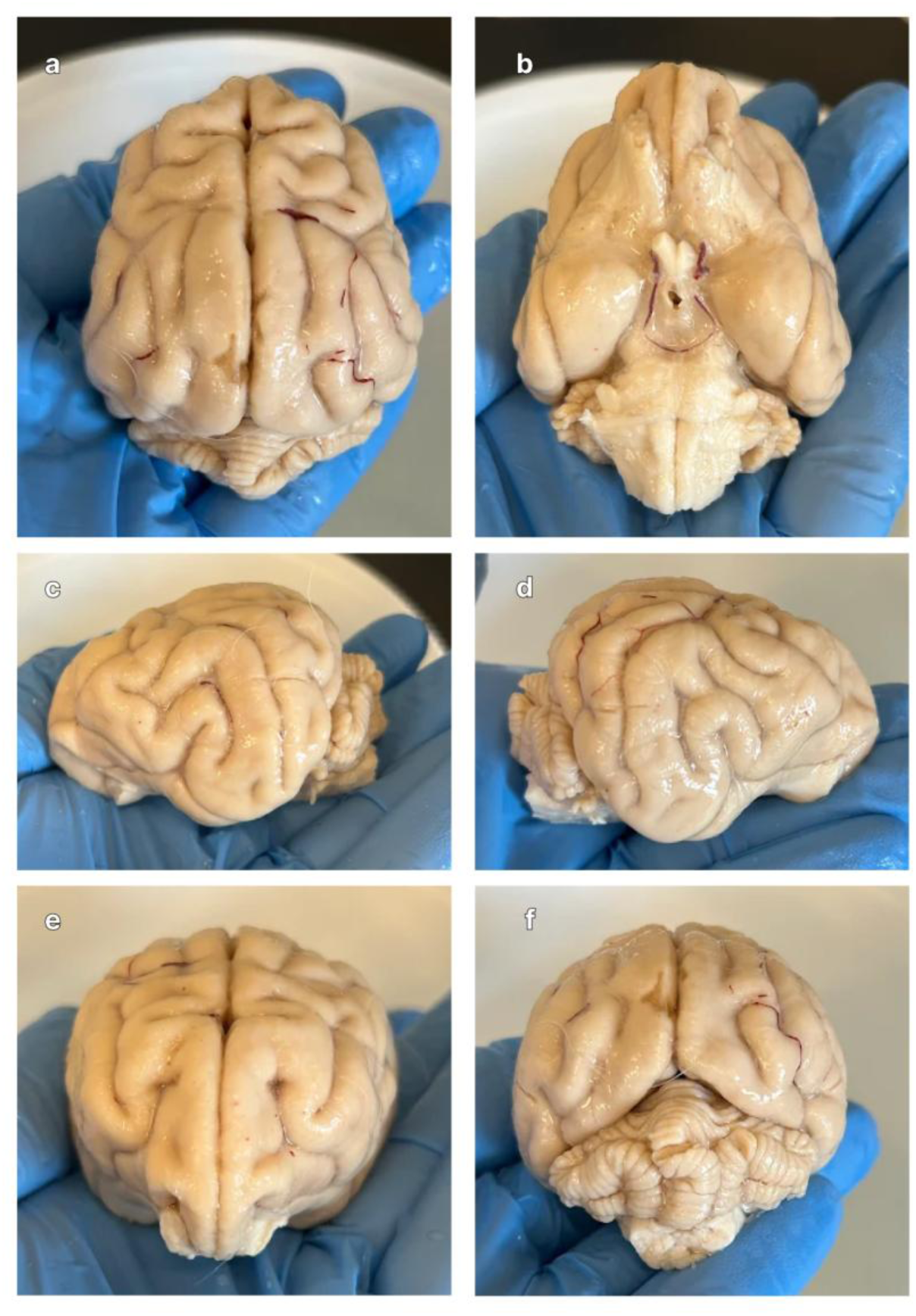
Gross appearance of donor 327 following perfusion with 20% neutral buffered formalin (NBF) containing 20% mannitol. Views are shown from the (**a**) superior, (**b**) inferior, (**c**) left lateral, (**d**) right lateral, (**e**) anterior, and (**f**) posterior aspects.

**Figure 5.**
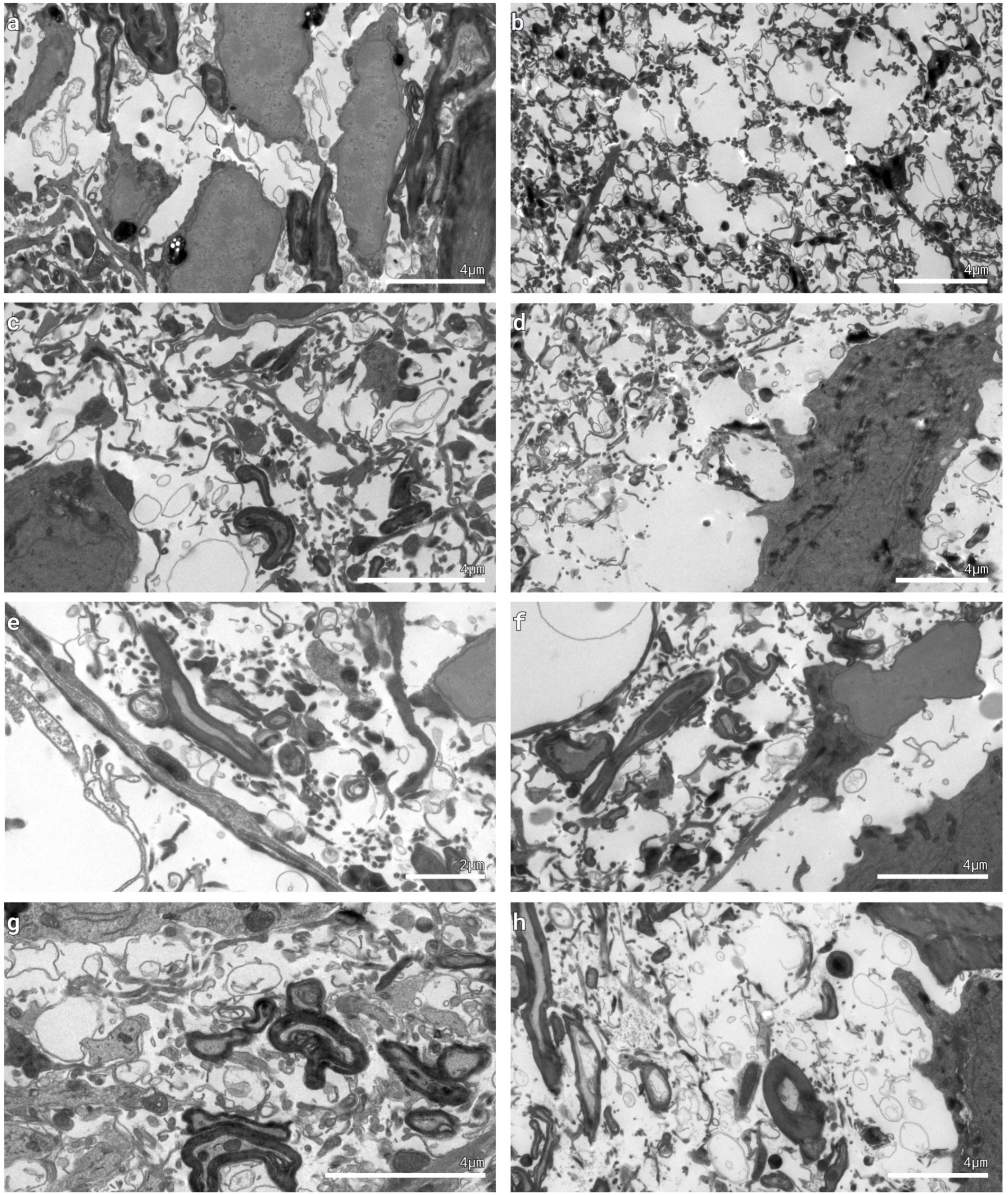
Representative electron micrographs from canine donor 327 following perfusion with 20% neutral buffered formalin containing 20% mannitol. Images are shown from the (**a**) left cerebellum, (**b**) right cerebellum, (**c**) left frontal cortex, (**d**) right frontal cortex, (**e**) left temporal cortex, (**f**) right temporal cortex, (**g**) thalamus, and (**h**) temporal white matter. Scale bars: all 4 µm, except (**e**) 2 µm. Images were acquired using a HT7500 transmission electron microscope using an AMT NanoSprint12 12-megapixel CMOS TEM Camera System.

### Comparison to previous cases perfused without dehydrating additives

We have previously reported perfusion fixation and electron microscopy results on brain tissue from human and canine donors perfused with NBF without dehydrating additives [19]. In these previous cases, electron microscopy revealed generally preserved ultrastructure with artifacts that are also commonly observed in immersion-fixed brain tissue, including swollen astrocytic processes and some extent of non-membrane-bound ambiguous interstitial zones [25,19,24]. The most notable perfusion-specific artifact we noticed in these cases was occasional vasogenic edema, which occurs when the blood-brain barrier has broken down so much that there is extravasation of the perfusate into the parenchyma [6].

When compared with these previously reported cases perfused with 10% NBF alone, the brains perfused with 10% mannitol and 10% PEG35 added to NBF generally showed substantially poorer ultrastructural preservation (**Figure 6**). Whereas the NBF-only cases demonstrated relatively clear cellular membranes and myelin architecture, the cases with hyperosmotic perfusate demonstrated more areas with expansion of the extracellular space, shrunken cellular processes, and distorted cell membranes.

**Figure 6.**
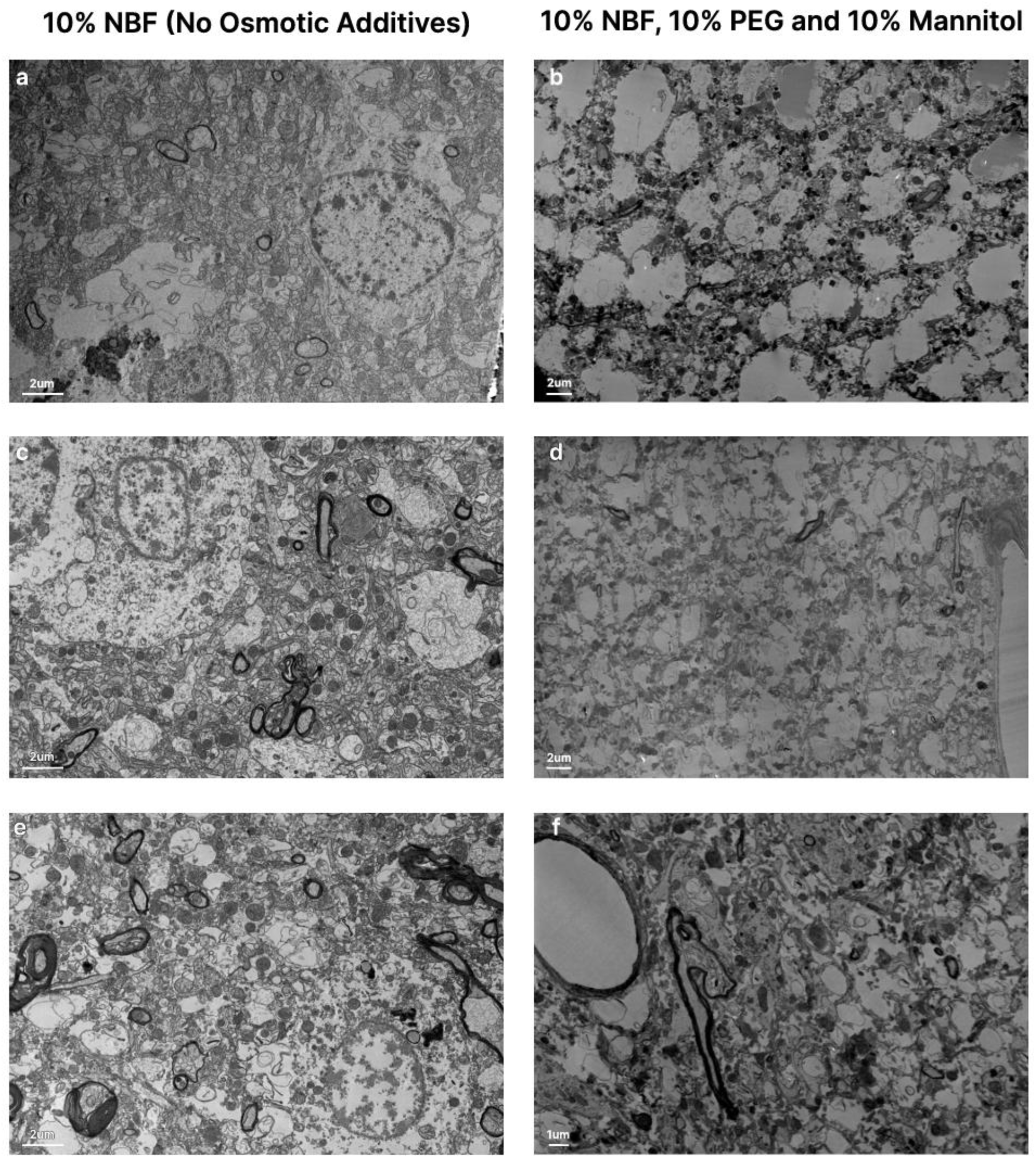
Comparison of electron microscopy following perfusion with standard and hyperosmotic perfusates. Left column (**a**, **c**, **e**): donors perfused with 10% neutral buffered formalin (NBF) alone. Right column (**b**, **d**, **f**): donors perfused with 10% neutral buffered formalin containing 10% mannitol and 10% PEG35. Canine donors are shown in (**c**) and (**d**); all other images are from human donors. The donor IDs are: 7 (**a**), 65 (**c**), and 34 (**e**) for the NBF group, and 247 (**b**), 250 (**d**), and 252 (**f**) for the hyperosmotic group. Scale bars: 2 µm in (**a-e**) and 1 µm in (**f**).

### Limitations

This study has several limitations. First, we were unable to perform a quantitative comparison of the perfusion quality between the hyperosmotic perfusate cases and our previously reported cases without osmotic additives because of the limited sample size. Second, although our findings support hyperosmotic dehydration as the cause of the observed ultrastructural abnormalities, we cannot completely exclude contributions from other protocol differences, such as differences in the perfusion pressure, or differences in donor characteristics, such as the duration of ischemia prior to preservation. Third, our sample size is small. Therefore, this manuscript should be considered a descriptive report of an artifact we observed in this series of cases, rather than an attempt to estimate the frequency of that artifact. Fourth, because electron microscopy samples only a very small fraction of the brain, we cannot determine the extent to which the ultrastructural artifacts we observed may have varied across unsampled regions.

Finally, we compared the ultrastructural abnormalities observed in brains perfused with 10% mannitol and 10% PEG35 as osmotic additives with those observed in a canine brain perfused with 20% mannitol as the sole osmotic additive. The similar abnormalities seen under both conditions suggest that severe hyperosmotic dehydration is sufficient to produce this osmotic shock artifact. However, our study was not designed to compare the relative effects of different osmotic additives.

## Conclusions

A pale, blood-free, and uniformly firm brain has long been considered a desirable endpoint indicating successful perfusion fixation [26]. Because osmotic additives have been proposed to improve perfusion quality following ischemia, we evaluated the use of perfusion with hyperosmotic fixative solutions in a small sample of human and canine donors. Gross, radiological, and histological evidence of vascular clearance indicated adequate to favorable perfusion by our standard assessment criteria in these cases. However, electron microscopy revealed a severe osmotic shock artifact in all sampled regions that had been perfused with these hyperosmotic solutions. Our study was not designed or powered to determine whether these hyperosmotic perfusates improve perfusate distribution relative to conventional perfusates, but our results do demonstrate that perfusion quality can be relatively high even when the ultrastructural preservation quality is severely compromised. We therefore suggest that studies developing perfusion protocols for connectomics should validate the ultrastructural preservation achieved with their perfusates directly rather than relying solely on macroscopic or radiological measures of perfusion quality.

## Author contributions

A.Sl., M.G., K.F., J.F.C., and A.T.M. conceptualized the article. A.Sl., M.G., K.H., A.So., W.J. performed electron microscopy experiments. E.L.T. and C.D.S. performed light microscopy experiments. A.Sl., M.G., K.H., and A.T.M. performed data analysis. A.T.M. wrote the initial draft of the manuscript. All authors reviewed the manuscript and all authors approved the final manuscript.

## Abbreviations

BBB: Blood-brain barrier
CT: Computed tomography
CV: Coefficient of variation
EM: Electron microscopy
H&E: Hematoxylin and eosin
MAID: Medical aid in dying
NBF: Neutral buffered formalin
NR: Not recorded
PBS: Phosphate-buffered saline
PEG: Polyethylene glycol
PEG35: Polyethylene glycol 35 kDa
PMI: Postmortem interval
STEM: Scanning transmission electron microscopy
TEM: Transmission electron microscopy

## Acknowledgements

Electron microscopy tissue preparation and imaging were performed at The Microscopy and Advanced Bioimaging CoRE at the Icahn School of Medicine at Mount Sinai. We would like to acknowledge the Neuropathology Brain Bank & Research CoRE at the Icahn School of Medicine at Mount Sinai for their histology and tissue processing services. The Icahn School of Medicine at Mount Sinai provided access to library resources.

## Funding

This work was supported by the Rainwater Charitable Foundation as well as NIH grants P30 AG066514, RF1 AG062348, K01 AG070326, and R01 NS146414. The funders had no role in the design of the study or in the collection or interpretation of the data.

## Conflict of interest

Andrew McKenzie, Andria Slaughter, Macy Garrood, and Katelyn Hedden are employees of Sparks Brain Preservation, a non-profit brain preservation organization.

## Data availability

The WSI and EM data are publicly available on Zenodo at the following DOIs: 10.5281/zenodo.21224468, 10.5281/zenodo.21224713, 10.5281/zenodo.21224581, 10.5281/zenodo.21224663, 10.5281/zenodo.21498531, and 10.5281/zenodo.21085501.

## Declaration of Generative AI Technologies

In the preparation of this manuscript, the authors used Claude (Anthropic) and ChatGPT (OpenAI) to improve the manuscript’s language. All AI tool-assisted content was reviewed and edited by the authors, who take full responsibility for the final publication.

